# Evolutionary analysis reveals repeated diversification events in immune metabolic pathways

**DOI:** 10.1101/2025.07.16.665195

**Authors:** Bruno W Fernandes Silva, Gleyson M Azevedo, Rodrigo JS Dalmolin

## Abstract

The human immune system is a complex, multifunctional network essential for host defense, tumor surveillance, and tissue repair. While conventionally divided into rapid-response innate immunity and antigen-specific adaptive immunity with memory, both modules operate synergistically through dynamic metabolic interactions that fuel immune responses. Although host-pathogen coevolution is recognized as a major evolutionary driver, the establishment scenario of immune metabolic pathways remains poorly characterized. Crucially, a systematic understanding of how the emergence of vertebrate malignancy can influence immune adaptations is needed. We analyzed 1,063 genes from 21 KEGG Pathway immune metabolic pathways and 1,124 cancer-associated genes from OncoKB/COSMIC. Evolutionary rooting was performed using the R package GeneBridge, inferring the most probable origins for each Cluster of Orthologous Genes (COG) across a 476-species eukaryotic phylogeny. Four clades showed significant immune orthologous groups (OGs) emergence: Metamonada, SAR, Choanoflagellata, and Actinopterygii. While cancer OGs diversified primarily during multicellular organism origins, immune OGs exhibited multiple diversification peaks, most prominently during jawed vertebrate emergence. Our findings demonstrate that human immune metabolic pathways underwent recurrent adaptive events during evolution, with marked complexity escalation in jawed vertebrates. We propose that malignant neoplasm emergence, coupled with epithelial-immune coevolution, served as complementary selective pressure driving progressive refinement of vertebrate immune mechanisms.

## Introduction

The human immune system comprises cells, molecules, organs, and fluids that function coordinately to defend the body against health threats, including infections and pathogens, while maintaining tumor surveillance and tissue repair capabilities (Sam-Yellowe 2021a; Poon and Farber 2020). Its core components are leukocytes (white blood cells), divided into granulocytes (neutrophils, eosinophils, basophils, mast cells) and agranulocytes (monocytes, macrophages, lymphocytes, and dendritic cells) (Shilts et al. 2022; Rich and Chaplin 2019). Their orthologous genes have been identified in distant taxa like sponges and cnidarians, participating in diverse biological processes including growth, development, cellular signaling, and stress responses (Ni et al. 2024; Williams and Gilmore 2020; Shivers et al. 2010).

Given the system’s complexity, it is conventionally divided into two major modules: (1) innate immunity, providing rapid, nonspecific defense, and (2) adaptive immunity, which mediates antigen-specific responses with immunological memory (Keselowsky et al. 2020; Rich and Chaplin 2019). These modules operate synergistically through continuous crosstalk with host metabolism, forming an integrated architecture enabling rapid and adaptable responses to environmental and physiological challenges (Buck et al. 2017; Odegaard and Chawla 2013). Metabolic pathways are pivotal in this context, supplying energy and substrates for immune cell activation, differentiation, and function, while directly instructing cellular behavior to regulate processes from proliferation to inflammatory mediator production (O’Neill et al. 2016). Thus, studying the immune system through its associated metabolic pathways is essential for a functional understanding of its evolutionary establishment.

From a systems biology perspective, evolutionary analyses are powerful tools to decipher how natural selection and other evolutionary processes shaped system structure, function, and dynamics over time. Immune system evolution occurred in layered phases: primordial innate mechanisms (e.g., barriers, molecular pattern recognition, basic cellular responses) emerged in most multicellular organisms (Wein and Sorek 2022; Cooper and Herrin 2010), while jawed vertebrates developed adaptive immunity, featuring lymphocytes with highly diversified antigen receptors and memory capacity (Smith et al. 2019; Cooper and Alder 2006). Host-pathogen coevolution is considered a major driver of immune diversification, forging the system into a complex, adaptable defense network (Iwasaki and Medzhitov 2015; Danilova 2006).

Epithelial somatic renewal is linked with immune surveillance, particularly in barrier tissues like skin and gut (Biton et al. 2018). Continuous epithelial turnover relies on immune signals to balance stem cell proliferation and differentiation, maintaining barrier integrity and rapid injury/infection responses (Guenin-Mace et al. 2023; Shin et al. 2022). Notably, 80–90% of human cancers originate epithelially (carcinomas) (McCaffrey and Macara 2011). Loss of epithelial hallmarks (e.g., polarity, cell adhesion) and dysregulated stem cell renewal/differentiation are tumorigenic hallmarks linked to cancer progression (Ribatti et al. 2020; Thompson and Nagaraj 2018). While key oncogenic genes/pathways are evolutionarily conserved, metastatic cells first appear in jawless vertebrates (lampreys) (Robert 2010). Understanding the evolutionary relationship between immune diversification and malignant cell emergence may reveal the extent to which tumor surveillance acted as a selective pressure driving vertebrate immune complexity.

Our study aimed to: (1) reconstruct the evolutionary scenario of human immune pathway establishment, and (2) compare this pattern with OG diversification in cancer. We analyzed 1,063 protein-coding genes from 21 immune-related metabolic pathways (KEGG Pathway), alongside 1,124 cancer-associated protein-coding genes from OncoKB and COSMIC. Using (Castro et al. 2008)‘s methodology, we identified vertically inherited archetypes within each OG to trace evolutionary origins. Results revealed multiple significant OG emergence events, reflecting distinct metabolic pathway diversification phases. Comparative analysis revealed divergent patterns, with cancer-associated OGs diversifying primarily during the origins of multicellular organisms, specifically between choanoflagellates and ctenophores. Whereas immune OGs exhibited multiple diversification peaks, most prominently during jawed vertebrate emergence. These findings suggest that increasing structural/functional complexity (including epithelial renewal systems) likely demanded more sophisticated immunological adaptations.

## Materials and Methods

### Gene list selection

Human immune system genes were obtained from the KEGG Pathway Database (Kanehisa and Goto, 2000; Kanehisa et al., 2017). All metabolic pathways listed under the Immune System section, within the Organismal Systems category, were selected. The Toll and Imd signaling pathway (04624) was excluded, as it is a microbial infection response pathway specific to insects. This selection resulted in a total of 21 metabolic pathways, comprising 1,063 protein-coding genes distributed across 469 orthologous groups (Fig. 1).

**Fig. 1.**
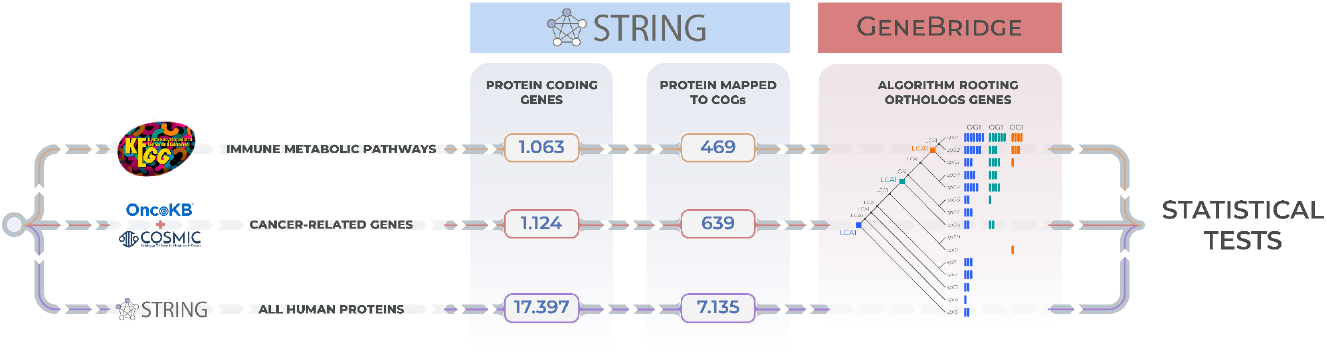
Data processing workflow for evolutionary root inference. Schematic representation of databases utilized and preprocessing steps enabling GeneBridge to infer the last common ancestor (LCA) with highest probability between the emergence of each orthologous group and *Homo sapiens*.

To construct a comprehensive cancer-associated gene list, we integrated data from two primary sources: OncoKB and the COSMIC database. OncoKB (https://www.oncokb.org/cancer-genes), a manually curated resource of oncogenic variants, provides clinically validated genes with therapeutic relevance (Chakravarty et al. 2017; Suehnholz et al. 2024). COSMIC (Catalogue Of Somatic Mutations In Cancer) supplemented this list with systematically curated genes recurrently mutated across tumor types (Bamford et al. 2004). These two sets were aggregated, forming a list with 1,124 protein-coding genes, comprising 639 groups of orthologs (Fig. 1).

The complete human protein-coding gene set was retrieved from the STRING database (https://string-db.org/cgi/download) (Szklarczyk et al. 2015). We selected *Homo sapiens* as the target organism and downloaded the compressed data file (9606.protein.info.v12.0.txt.gz) for subsequent filtering. This gene list has 17,397 protein-coding genes, which are part of 7,135 orthologous groups (Fig. 1).

### Evolutionary Rooting Analysis

To trace the history of immune gene emergence, an evolutionary rooting analysis was performed using GeneBridge (Campos et al. 2024), an R package developed for large-scale evolutionary gene analyses based on the distribution of OGs (Orthologous Groups). All genes contained annotations for at least one Cluster of Orthologous Groups (COG), meaning each COG could contain one or more genes. GeneBridge infers the most likely evolutionary root of an OG by analyzing the distribution of its genes across the branches of a reference phylogenetic tree. We used a 476-species eukaryotic tree from STRING. The tree was obtained via the R package AnnotationHub (Morgan M and Shepherd L, 2025), which provides an interface for accessing and managing genomic annotations and other public database resources.

### Statistical Tests

To identify clades with a significant increase of orthologs, the MAD-z-score (Median Absolute Deviation) was employed (Iglewicz and Hoaglin 1993). This metric is more suitable than the conventional z-score because orthologs emergence values vary considerably across different clades of the reference phylogenetic tree, making the median absolute deviation (MAD) more robust than the traditional standard deviation. Modified z-score was performed using orthologs emergence values per clade. The hypergeometric test was performed to identify the points where there was a significant emergence of orthologs from immune metabolic pathways in the clades of the reference phylogenetic tree. The values used for the test were the following: N = total of early immune gene set; n = number of orthologs within the clade; and M = number of genes associated with each metabolic pathway; k = orthologs in the clade also linked to the pathways. The p-value was derived from the hypergeometric distribution using the phyper function in R (v.4.4.2).

## Results

### Evolution of the human immune system

Our study selected genes involved in annotated metabolic pathways related to the immune system. In total, 1,063 protein-coding genes comprising 21 pathways from the KEGG database were analyzed, with orthology information mapped for 469 orthologous groups using STRING. The number of genes per metabolic pathway varied considerably, with the Chemokine signaling pathway, Neutrophil extracellular trap formation, and NOD-like receptor signaling pathway being the three pathways containing the highest number of genes (supplementary Fig. 1A). The vast majority of genes were annotated for only one immune metabolic pathway (754 genes), while a small subset was present in six or more pathways (supplementary Fig. 1B). Our analysis revealed that immune-metabolic pathways exhibit low molecular crosstalk between distinct biochemical routes.

The rooting analysis performed using the R package GeneBridge inferred the most recent common ancestor between each orthologous group comprising the gene list and *Homo sapiens*. This approach allowed us to determine how many OGs emerged in each clade of the reference tree. The test revealed four clades with a significant number of emerging OGs: Metamonada, SAR, Choanoflagellata, and Actinopterygii (Fig. 2). Another notable observation was the period of low orthologs acquisition between Ctenophora and Tunicata, coinciding with the emergence and extensive diversification of invertebrate organisms. Despite representing a substantial evolutionary timeframe encompassing remarkable organismal diversity, this interval appears to have contributed minimally to the repertoire of orthologs subsequently conserved in humans.

**Fig. 2.**
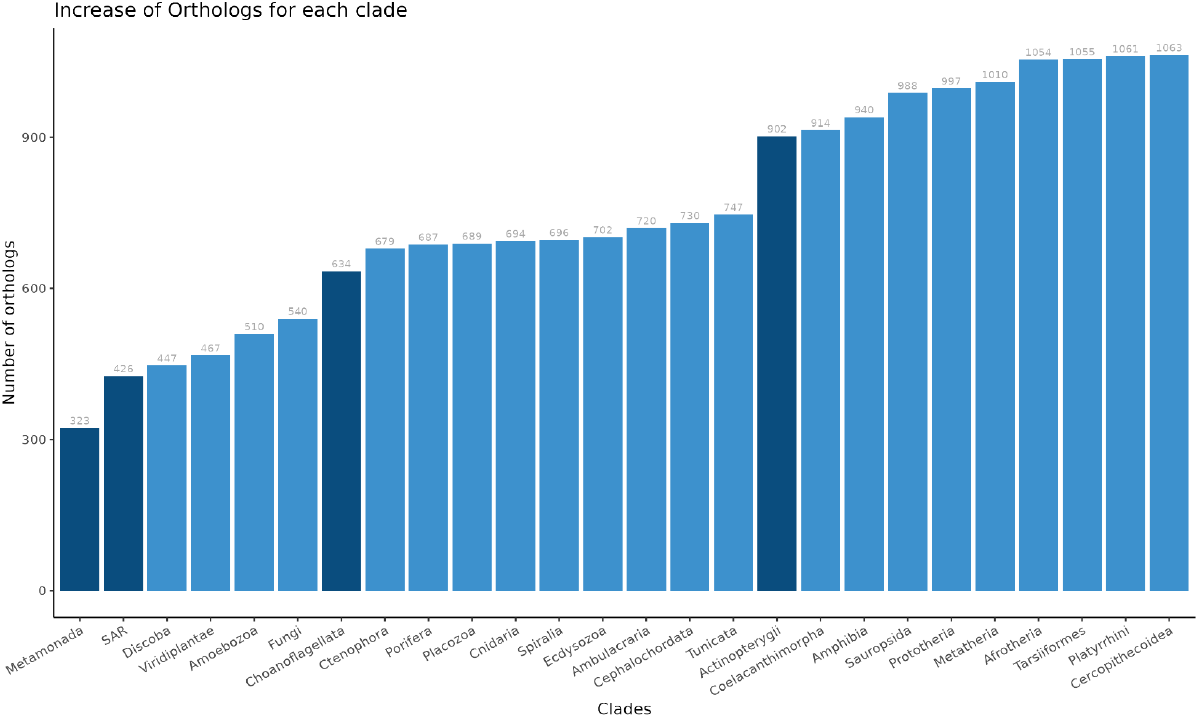
Orthologs emergence across evolutionary clades. The chart is organized chronologically from ancestral (left) to derived (right) clades, with orthologs counts incrementally adjusted by the number of emerging orthologs per clade. Modified z-score was performed using orthologs emergence values per clade to identify orthologs emergence events. The highlighted clades represent the largest orthologs emergences; the threshold used was (p-value < 0.05).

### Identification of diversification events in immune metabolic pathways

The hypergeometric test was performed to identify diversification events of immune metabolic pathways across evolutionary clades where orthologs were rooted. For most pathways, multiple diversification events were identified (Fig. 3). In Metamonada, five metabolic pathways exhibited significant orthologs emergence: Neutrophil extracellular trap formation, T cell receptor signaling pathway, B cell receptor signaling pathway, Platelet activation, and C-type lectin. SAR, one of the clades with the highest gene emergence, the highlighted pathways were Platelet activation and Fc gamma R-mediated phagocytosis. Choanoflagellata displayed three prominent pathways: Platelet activation, Chemokine signaling pathway, and Fc epsilon RI signaling pathway. Notably, three of the four clades with the highest orthologs emergence showed significant expansion in Platelet activation. The clade with the greatest number of enriched pathways was Actinopterygii, totaling six: Hematopoietic cell lineage, IL-17 signaling pathway, Th17 cell differentiation, Th1 and Th2 cell differentiation, Antigen processing and presentation, and Intestinal immune network for IgA production. Two metabolic pathways exhibited particularly notable evolutionary patterns, each demonstrating five significant diversification events: Th17 cell differentiation and the complement and coagulation cascades, with emphasis on the latter due to the intervals where the events occurred, where the first point of emergence occurred in Discoba and the most recent in Cercopithecidae, the most derived clade where GeneBridge inferred immune orthologs emergence.

**Fig. 3.**
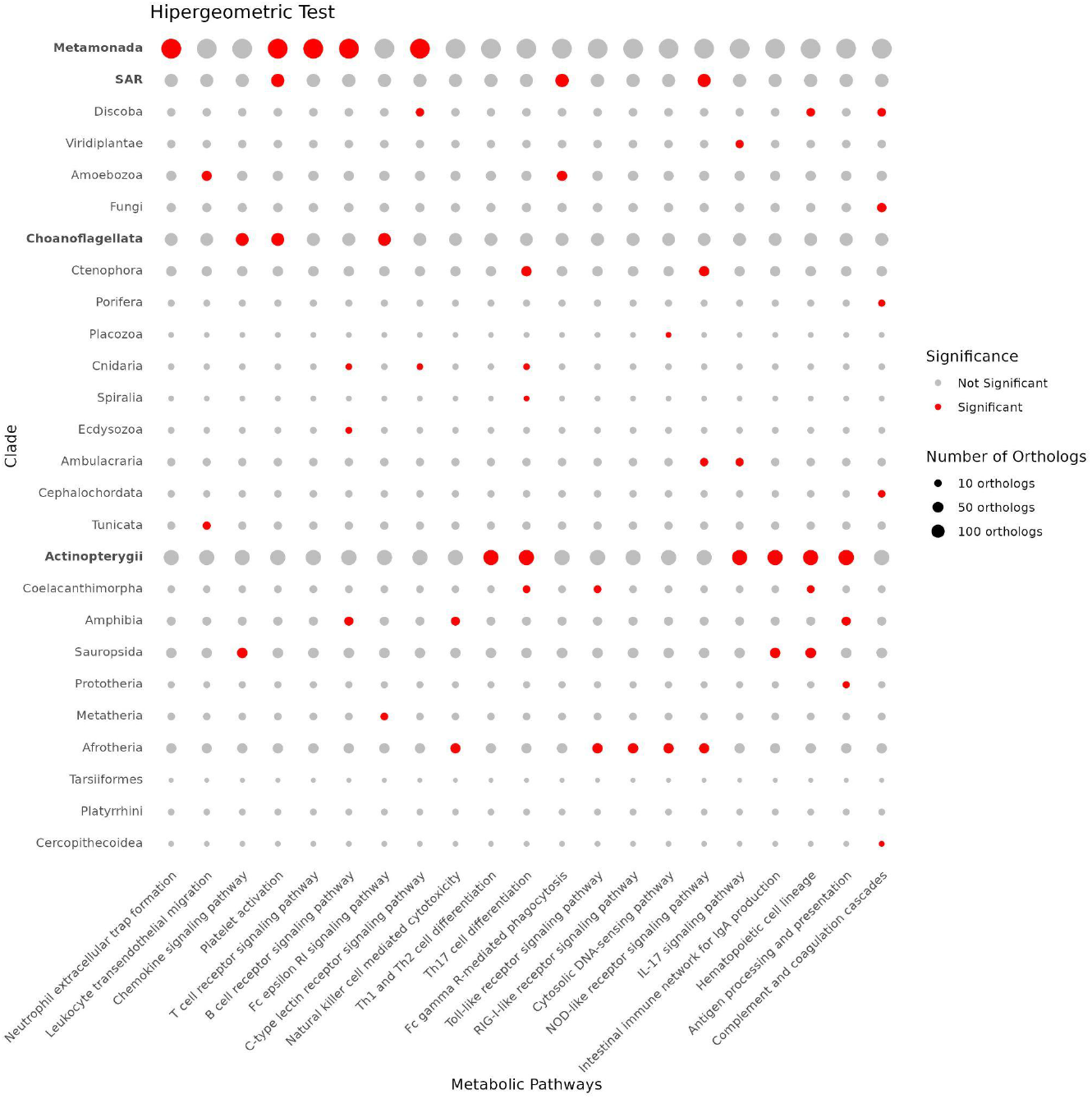
Significant orthologs emergence events across metabolic pathways. Dot plot showing results of hypergeometric testing, highlighting clades with statistically significant orthologs emergence for each immune-related metabolic pathway analyzed. Clades are ordered from ancestral (top) to recent (bottom). Red points indicate significant emergence of orthologs from immune metabolic pathways in clades (p-value < 0.05).

### Comparing the immune evolutionary scenario with cancer OGs emergence

Our analyses revealed similar clades with a greater increase in orthologs among immune and cancer-related OGs (Fig. 4). Evolutionary reconstruction demonstrated that cancer-related OGs underwent a major diversification event coinciding with the origin of multicellular organisms, between Choanoflagellata and Ctenophora. Although immune OGs have also diversified significantly in Choanoflagellata, they do not follow the trend demonstrated by cancer-related orthologs in Ctenophora. The largest diversification event of OGs involved in immune metabolic pathways occurred in Actinopterygii, with an increase of approximately 15% in new OGs.

**Fig. 4.**
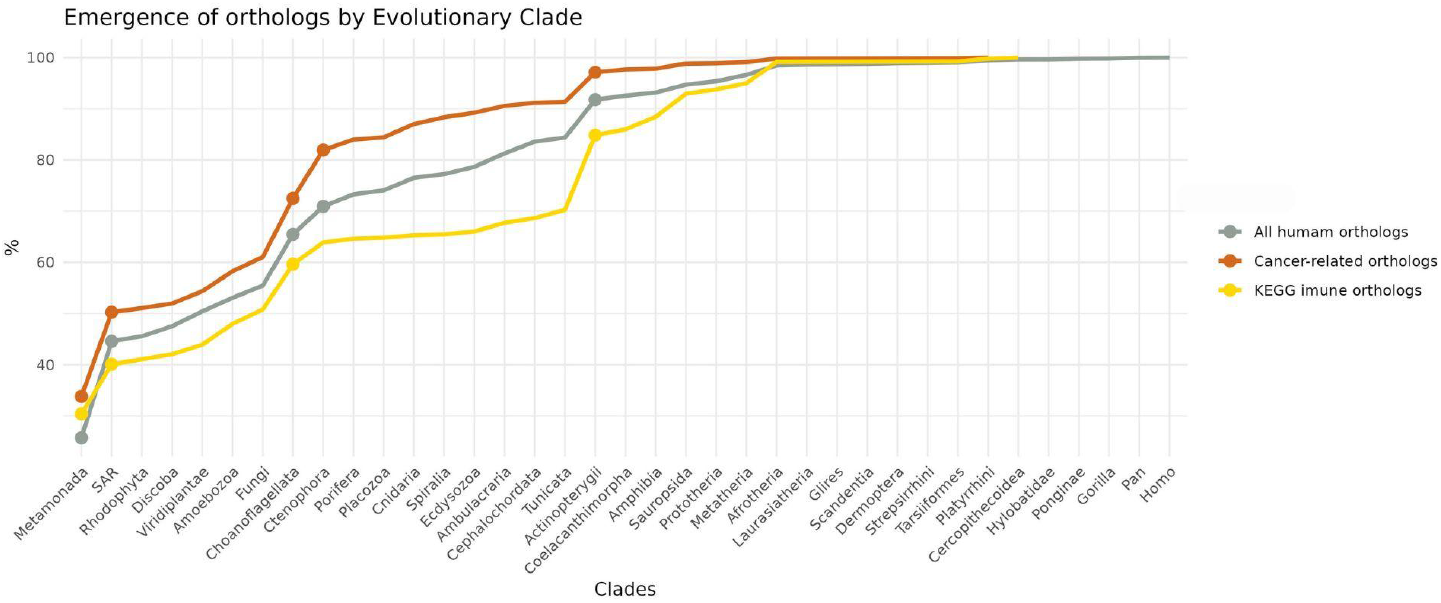
Comparative evolutionary emergence of immune and cancer-related OGs. The line plot displays normalized OG emergence timelines, with the gray line representing all Homo sapiens OGs (control). Immune-related (yellow) and cancer-related (orange) OGs are shown for comparison, highlighting their distinct evolutionary trajectories. Circles denote the significant emergence of OGs, defined by a modified z-score > 3.5 and p-values < 0.05.

As a control, we performed evolutionary rooting of all human OGs. This comparative analysis revealed that immune metabolic pathways exhibit a distinct pattern from the general human protein pool: while the curve for all human OGs follows a similar pattern to those related to cancer (between Choanoflagellata and Ctenophora), followed by steady gains throughout invertebrate evolution (up to Tunicata), immune OGs exhibited minimal increases during this latter period. Thus, the origin of vertebrates, represented by the Actinopterygii clade in our analysis, appears to represent a crucial evolutionary landmark for the diversification of the immune system.

## Discussion

For decades, evolutionary studies have sought to understand how the biological defenses comprising the human immune system emerged, diversified, and adapted over time (Liston et al. 2021). These investigations integrate genetics, comparative biology, model organism experiments, and other approaches to elucidate immune system evolution and its adaptive maintenance (Jiao et al. 2024; Vinkler et al. 2023; Schulenburg et al. 2004). Critical gaps remain regarding the evolutionary scenario in which the diversification and establishment of the immune system occurred. Within this context, our study proposes a systemic approach to unravel the evolutionary scenario of immune system establishment, focusing on metabolic pathway analysis as an investigative tool. Our results demonstrate that, quantified by orthologous group (OG) numbers, immune system diversification occurred predominantly across four key evolutionary clades: Metamonada, SAR, Choanoflagellata, and Actinopterygii. This OG emergence pattern aligns logically with both gene quantity and system complexity, suggesting multiple adaptive events throughout evolutionary history (Subramanian et al. 2015).

The Metamonada clade comprises three orders: Retortamonadida, Diplomonadida, and Oxymonadida, forming a supergroup of anaerobic, flagellated unicellular organisms (Brugerolle 1991). As the most basal clade in the reference phylogenetic tree used by the GeneBridge algorithm, its high number of emerging OGs also reflects prokaryotic-derived immune modules conserved in eukaryotes (Bernheim et al. 2024; Wein and Sorek 2022). Notably, approximately 51% of orthologs in our immune gene list had their last common ancestors inferred to predate Choanoflagellata, suggesting that most proteins participating in immune metabolic pathways have ancient evolutionary origins rooted in basal clades. For instance, several gene families rooted in early-diverging clades (e.g., calmodulin CALM, glycogen synthase kinase GSK, phospholipase C PLC, Nuclear Factor kappa B NF-κB, Myosin Light Chains MYL, mitogen-activated protein kinase MAPK, and the phosphatase proteins PPP1, PPP2, and PPP3) are involved in non-immune-exclusive cellular processes such as proliferation, signaling, and cell cycle regulation (Xu et al. 2017; Perkins 2012; Uhrig et al. 2013; Saidi et al. 2012).

Our results showed complex pathways such as T-cell receptor (TCR) signaling and neutrophil extracellular trap (NET) formation, with only one diversification event located in Metamonada. NETs are web-like structures released by neutrophils that play essential roles in defense against infections. They capture and eliminate microorganisms but are also involved in inflammatory response regulation (Wang et al. 2024; Islam and Takeyama 2023). Analogous extracellular trap mechanisms have been observed in some invertebrates, but classic NETs and TCR signaling processes as seen in humans appear to be exclusive to jawed vertebrates (Díaz-Godínez and Carrero 2019; Danilova 2006). These Metamonada results suggest that proteins rooted in more basal periods may have been co-opted for functions in more derived pathways in evolutionarily recent clades.

Choanoflagellata exhibit high emergence of new OGs, representing ∼9% of the total gene list, primarily associated with the chemokine signaling pathway, a system essential for cellular migration in multicellular organisms (Kohli et al. 2022). Recent research revealed that *Monosiga brevicollis* (a choanoflagellate species) possesses the STING gene, known in animals for activating innate immune pathways in response to pathogen-derived molecular signals (Woznica et al. 2021). Furthermore, earlier studies demonstrated that holozoans like choanoflagellates can transition between cellular states using mechanisms homologous to metazoans, including transcription factors, enhancers, promoters, and non-coding RNAs (Sebé-Pedrós et al. 2017; Gaiti et al. 2017; Sebé-Pedrós et al. 2016). These findings imply that choanoflagellates already harbored molecular machinery precursor to vertebrate immune functions, such as cellular migration and specification, suggesting that key immunological complexity predated metazoan radiation

Our analyses revealed low emergence of new OGs in invertebrates. Although these organisms lack vertebrate-like adaptive immunity, they exhibit sophisticated defense mechanisms encompassing physical barriers, cellular and molecular responses, and even pathogen-specific behavioral adaptations (Nakad et al. 2016; Loker et al. 2004). Recent studies demonstrate that invertebrates can develop immune responses with memory-like features and specificity, as observed in crustaceans and insects, where prior pathogen exposure enhances protection and enables maternal transfer of specific immunity (Kloc et al. 2024; Little et al. 2003). Furthermore, the molecular diversity of immune proteins facilitates targeted responses against distinct pathogen types, implying specialized recognition and defense capacities (Cerenius and Söderhäll 2013). In this context, the low OGs emergence does not indicate stalled evolutionary complexity in immune metabolic pathways. Rather, this period likely represented functional diversification through modifications of pre-existing pathways.

The origin of the adaptive immune system, characterized by the emergence of B and T lymphocytes and the major histocompatibility complex (MHC), is closely linked to the irradiation of jawed vertebrates (Ohta et al. 2019; Anderson et al. 2004). Our GeneBridge-derived results support this correlation by identifying in Actinopterygii the emergence of OGs associated with metabolic pathways essential for adaptive immunity, including: (i) antigen processing and presentation, and (ii) the Intestinal immune network for IgA production. Furthermore, we detected expansion of OGs related to Th1, Th2, and Th17 cell differentiation. These CD4+ helper T cell subsets specialize adaptive immune responses, enabling targeted reactions against distinct threat categories (Zhu and Zhu 2020). Each subtype activates unique mechanisms, promoting pathogen-specific responses against intracellular pathogens (Th1), parasites (Th2), or extracellular microbes (Th17) (Zhu and Zhu 2020; Kaymaz and Beikler 2019). These findings suggest that vertebrate emergence constituted a critical evolutionary transition for diversifying adaptive immunity mechanisms, particularly in immune response specialization.

One of the most critical innovations in vertebrates is somatic epithelial renewal, particularly in barrier tissues like skin and intestine. This continuous turnover mechanism, regulated through immune-derived signals (Guenin-Macé, Konieczny, and Naik 2023; Shin et al. 2022), orchestrates cellular proliferation and differentiation. For instance, helper T lymphocytes and innate cells secrete cytokines, such as interleukins and Transforming Growth Factor (TGF-β), that directly modulate epithelial stem cell fate, promoting differentiation during inflammation or self-renewal in homeostasis (Guenin-Macé, Konieczny, and Naik 2023). Nearly all nucleated cells express MHC class I, while MHC class II is typically restricted to antigen-presenting cells but can be induced in other cell types under certain conditions, expanding the immune system’s capacity to recognize and respond to threats (Sam-Yellowe 2021b). Recent studies have demonstrated that high expression of these molecules exhibits protective effects against most cancer types.(Schaafsma et al. 2021)

Given that most human cancers originate epithelially (carcinomas) (Ribatti et al. 2020; McCaffrey and Macara 2011), dysregulation of immune-mediated epithelial renewal emerges as a pivotal oncogenic driver (Guenin-Mace et al. 2023). Intriguingly, the earliest evolutionary records of malignant neoplasms appear in jawless vertebrates (lampreys) (Robert 2010), predating the period where our results demonstrate significant expansion of antigen processing/presentation pathways in jawed vertebrates. This temporal parallel suggests a coevolutionary scenario: (1) escalating epithelial renewal complexity and (2) malignant phenotype emergence acted as complementary selective pressures, leading to a progressive refinement of vertebrate immune surveillance mechanisms. That said, epithelial-immune interaction appears to represent a crucial adaptive landmark in the evolution of the immune system.

This study used a systems approach to investigate the human immune system, aiming to identify the evolutionary period at which these metabolic pathways became functionally integrated, as observed in more derived organisms. In summary, our results demonstrate that metabolic pathways associated with the human immune system have undergone multiple adaptive events throughout evolution, as evidenced by the pattern of emergence of OGs. Although over half of the analyzed OGs originated before metazoan diversification, our results suggest that the emergence of malignant neoplasms, coupled with the coevolution of epithelial-immune interactions, may have acted as important selective pressures for the diversification of vertebrate immune system. The selection of representative genes for biological systems poses considerable methodological challenges. This difficulty extends beyond the sheer number of genetic elements involved to encompass the pleiotropic nature of these genes, which frequently participate in multiple essential cellular processes, including diverse signal transduction pathways. While the GeneBridge approach enables reconstruction of the most probable last common ancestor through distributional analysis of orthologs, the methodology has inherent limitations in temporal precision when determining the exact origin of specific genes within each cluster. These considerations highlight the need for complementary studies to achieve technical refinement and consolidate the findings described in this work.

