## Supplementary figure 1 for "Evolutionary analysis reveals repeated diversification events in immune metabolic pathways"

### Supplementary Material

**A**

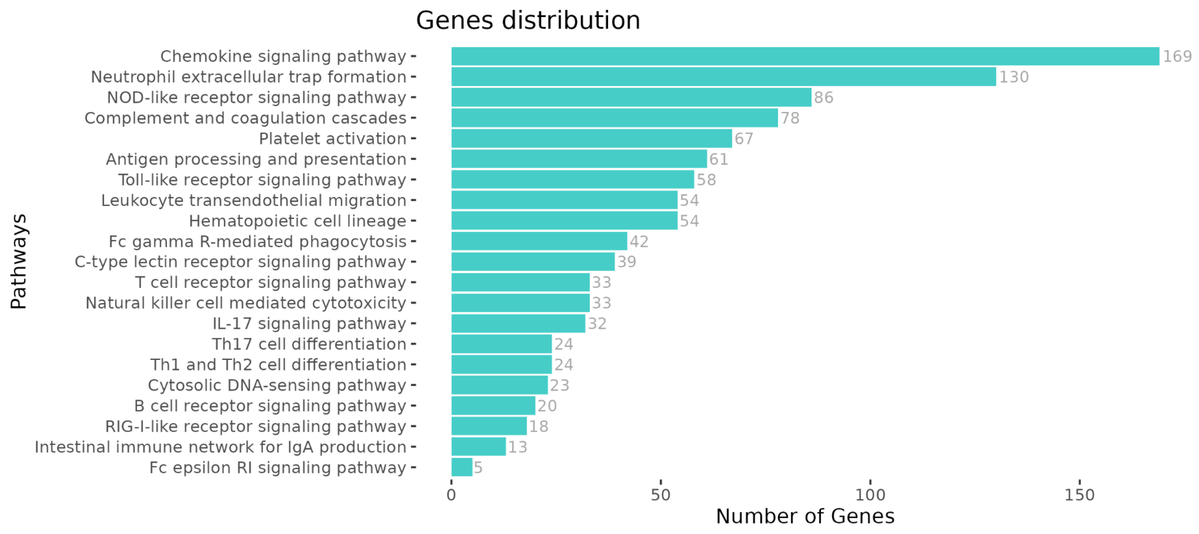


**B**

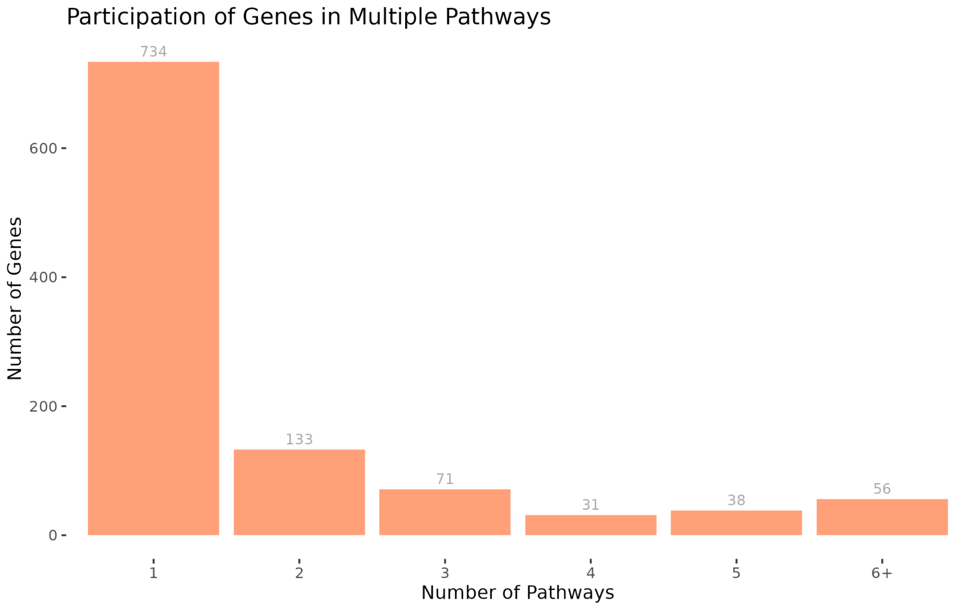


Fig. 1 - **Characterization of the immune gene dataset.** (A) Bar plot showing gene counts per KEGG immune metabolic pathway. (B) Gene distribution by pathway participation frequency: x-axis indicates number of associated pathways per gene, y-axis shows gene counts.
